# S1P controls endothelial sphingolipid homeostasis via ORMDL

**DOI:** 10.1101/2021.10.22.465503

**Authors:** Linda Sasset, Kamrul H. Chowdhury, Onorina L. Manzo, Luisa Rubinelli, Csaba Konrad, J. Alan Maschek, Giovanni Manfredi, William L. Holland, Annarita Di Lorenzo

**Affiliations:** Department of Pathology and Laboratory Medicine, Cardiovascular Research Institute, Weill Cornell Medicine, 1300 York Avenue, New York, NY, 10021; Brain and Mind Research Institute, Weill Cornell Medicine, 1300 York Avenue, New York, NY, 10021; Department of Nutrition and Integrative Physiology, University of Utah College of Health, Salt Lake City, UT, 84112; Department of Pharmacy, University of Naples “Federico II”, Naples, Italy

## Abstract

Sphingolipids (SL) are both membrane building blocks and potent signaling molecules regulating a variety of cellular functions in both physiological and pathological conditions. Under normal physiology, sphingolipid levels are tightly regulated, whereas disruption of sphingolipid homeostasis and signaling has been implicated in diabetes, cancer, cardiovascular and autoimmune diseases. Yet, mechanisms governing cellular sensing of SL, and according regulation of their biosynthesis remain largely unknown.

In yeast, serine palmitoyltransferase (SPT), catalyzing the first and rate limiting step of sphingolipid *de novo* biosynthesis, is negatively regulated by Orosomucoid 1 and 2 (Orm) proteins. Lowering sphingolipid levels triggers Orms phosphorylation, resulting in the removal of the inhibitory brake on SPT to enhance sphingolipid *de novo* biosynthesis. However, mammalian orthologs ORMDLs lack the N-terminus hosting the phosphosites. Thus, which sphingolipid(s) are sensed by the cells, and mechanisms of homeostasis remain largely unknown. This study is aimed at filling this knowledge gap.

Here, we identify sphingosine-1-phosphate (S1P) as the key sphingolipid sensed by endothelial cells via S1PRs. The increase of S1P-S1PR signaling stabilizes ORMDLs, which downregulates SPT activity to maintain SL homeostasis. These findings reveal the S1PR/ORMDLs axis as the sensor-effector unit regulating SPT activity accordingly. Mechanistically, the hydroxylation of ORMDLs at Pro137 allows a constitutive degradation of ORMDLs via ubiquitin-proteasome pathway, therefore preserving SPT activity at steady state. The disruption of the S1PR/ORMDL axis results in ceramide accrual, mitochondrial dysfunction, and impaired signal transduction, all leading to endothelial dysfunction, which is an early event in the onset of cardio- and cerebrovascular diseases.

The disruption of S1P-ORMDL-SPT signaling may be implicated in the pathogenesis of conditions such as diabetes, cancer, cardiometabolic disorders, and neurodegeneration, all characterized by deranged sphingolipid metabolism. Our discovery may provide the molecular basis for a therapeutic intervention to restore sphingolipid homeostasis.

## INTRODUCTION

Formed by the subunits SPT Long Chain 1 and 2 (SPTLC1 and SPTLC2), mammalian SPT activity is enhanced by small subunits ssSPTa and ssSPTb^1^, and decreased by ORMDLs^2^ and Nogo-B^3^. The requirement of sphingolipid *de novo* biosynthesis for viability and health is underlined by genetic evidence in humans and mice. SPTLC1/2 mutations cause Hereditary Sensory Neuropathy Type I^4,5^, SNPs in ORMDLs are associated to asthma^6^ and atherosclerosis^7^, while the excision of Sptlc1 or Sptlc2 genes in mice is embryonically lethal^8^. In yeast, Orms (Orm1 and Orm2) proteins regulate SL homeostasis, with the phosphorylation of Orms releasing the brake on SPT^2^. However, mammalian ortholog ORMDLs lack the N-terminal regions hosting these phosphosites^9^. How cells sense SL, monitor the rate of the *de novo* biosynthesis, and what goes awry in disease remain unknown.

S1P signaling is critical in development, physiological homeostasis, and diseases^10^. Genetic disruption of the S1P pathway results in congenital defects in humans, including Sjogren-Larsson syndrome, adrenal insufficiency and nephrosis, hearing impairment, embryonic lethality, and post-natal organ defects in mice, underlining that functional S1P signaling is a prerequisite for health. Within the cardiovascular and immune systems, S1P is necessary for vascular development^11^ and homeostasis^12^ as well as immune cell trafficking^13^, mainly via S1PR1. Endothelial S1PR1 controls blood flow and pressure via nitric oxide (NO) formation^3,14,15^, and maintains the quiescent state of the endothelium by exerting anti-inflammatory^16,17^ and barrier^18^ functions. The endothelium is also an important source of plasma ceramide^19^ and S1P^20^, which is transported outside of the cells by the bonafide transporter for S1P, Spinster-2 (Spns2)^21^. Autocrine S1P signaling controls flow-induced vasodilation, which is a vital function of blood vessels to meet the tissue metabolic demands^22^. Disruption of endothelial S1P signaling by deletion of S1PR1^15^ or Spns2^23^ results in vascular^15,23^ and barrier^18^ dysfunctions, severe hypertension^15,23^ and atherosclerosis^16^, underlying the fundamental role of S1P-S1PR1 signaling in preserving vascular health.

## RESULTS

### S1P inhibits SPT activity via ORMDLs stabilization

This study tested the hypothesis that S1P signaling is a fundamental molecular mechanism used by the cells to maintain sphingolipid homeostasis. The capability of S1PR to transduce S1P levels in a biological signal, and the specificity and dynamicity of this interaction, make of S1P-S1PR an ideal metabolite-sensor system to regulate cellular sphingolipid homeostasis through an effector yet to be identified. To test this hypothesis, we used murine EC (mEC) inducible knockout for *S1pr1* (**Fig. 1A**). Interestingly, *S1pr1* deletion resulted in increased SPT activity and SL levels (**Fig. 1A-C** and **Supplementary Fig. 1**) suggesting that S1PR1 signaling could function as negative feedback on sphingolipid metabolism. Based on this finding, we hypothesized that S1P is the key sphingolipid metabolite sensed by the cells to modulate the sphingolipid *de novo* biosynthesis via SPT (effector). Interestingly, S1P was able to rapidly decrease SPT activity of human umbilical vein EC (HUVEC, **Fig. 1D**). Recent studies suggested that ceramide could modulate the ORMDL-SPT complex^24–26^. In line with this finding, the addition of C16:0-ceramide significantly decreased SPT activity in HUVEC (**Fig. 1E**). However, this effect was abolished by SKI II, an inhibitor of sphingosine kinase-1 (SphK1) and SphK2 (**Fig. 1F**), which give rise to S1P via the phosphorylation of sphingosine, suggesting that S1P derived from ceramide metabolism mediates the inhibitory effects of ceramide on SPT. mEC with genetic deletion of *Sphk1* and *Sphk2* (Sphk1,2*^ECKO^*, **Fig. 1G**) corroborated that S1P formation is necessary for ceramide to downregulate SPT activity (**Fig. 1H**). Of note, basal SPT activity was significantly upregulated in Sphk1,2*^ECKO^* (**Fig. 1H**), revealing a direct role of endothelial-derived S1P in constraining SPT activity. Contrary to Orms in yeast, ORMDLs are not phosphorylated^24^ in mammals, and our data corroborated this finding (**Supplementary Fig. 2**). Interestingly, S1P stimulation triggered a rapid increase in ORMDLs levels (ca. 2.5-fold), without affecting Nogo-B, SPTLC1 or SPTLC2 expression (**Fig. 1I,L**). Ceramide also induced a rapid increase in ORMDLs (**Fig. 1M**), which was abolished by SKI II inhibitor, consistent with S1P mediating ceramide inhibition of SPT activity (**Fig. 1F**). Both NOGO-B (**Fig. 1N**) and ORMDLs (**Fig. 1O**) knock-down resulted in a higher SPT activity, in line with a constitutive inhibition of SPT. However, only knock-down of ORMDLs (**Fig. 1O**) abolished the S1P downregulation of SPT activity, suggesting that ORMDLs and not Nogo-B are accountable for a nimble regulation of SPT activity by S1P (**Fig. 1P**).

**Figure 1.**
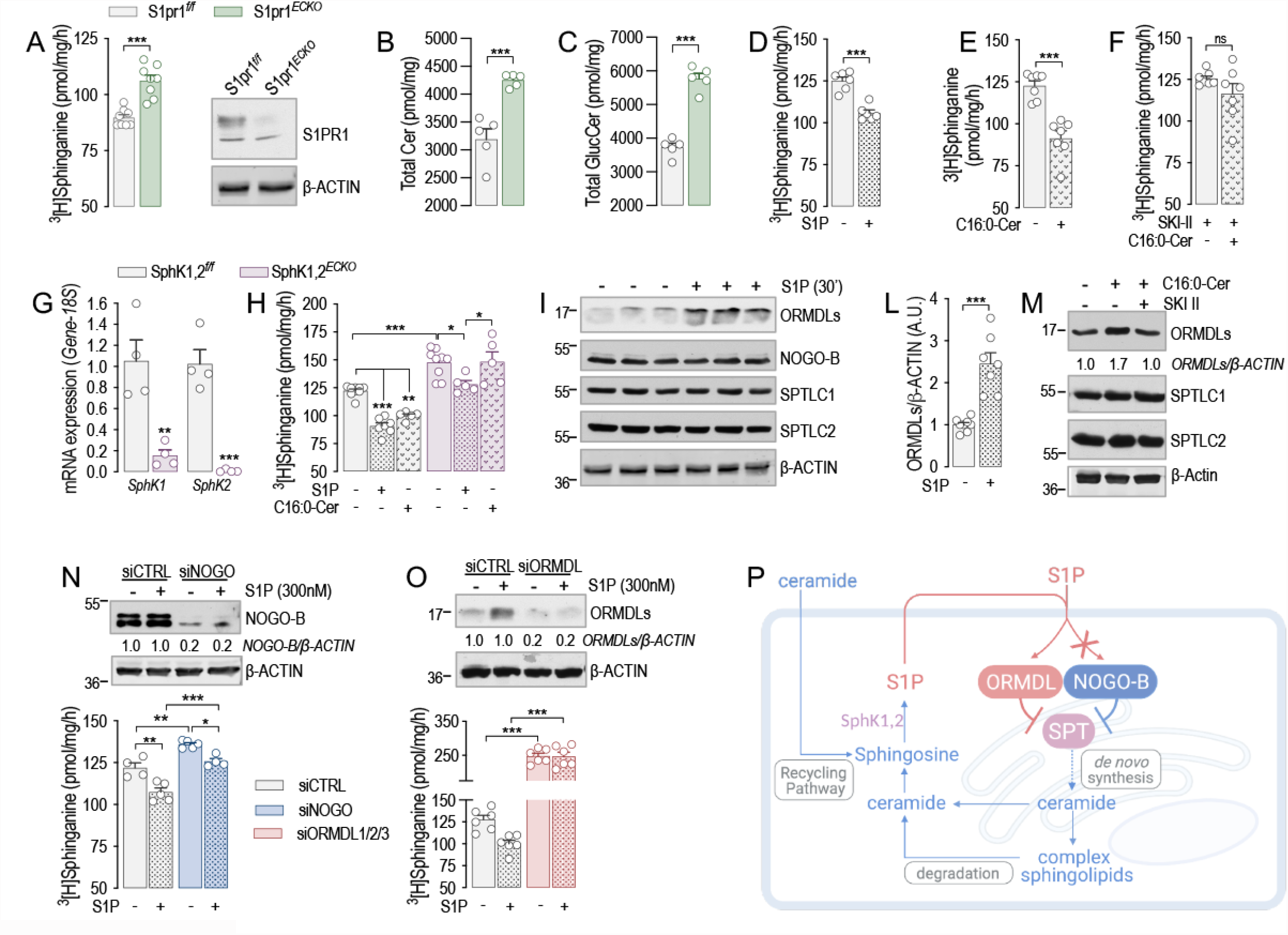
S1P inhibits SPT activity via ORMDLs stabilization. **(A)** SPT activity and Western blot (WB) analysis of S1PR1 in S1pr1*^f/f^* and S1pr1*^ECKO^* endothelial cells after 4-OHT (1μM, 72h) treatment. (n=8/group from 3 independent EC isolations/group; 4 mice/EC isolation). LC-MS/MS quantification of Total **(B)** Ceramide and **(C)** Glucosylceramide in S1pr1*^f/f^* and S1Pr1*^ECKO^* endothelial cells after 4-OHT (1μM, 72h) treatment (n=5/group from 2 independent EC isolations/group; 4 mice/EC isolation). SPT activity in HUVEC in **(D)** absence or presence of S1P (300nM, 30’) (n=6); **(E)** absence of presence of C16:0-ceramide (300nM, 30’) and **(F)** absence of presence of C16:0-ceramide (300nM, 30’), in presence of SKI II (1µM, 1h pre-treatment). **(G)** RT-PCR for SphK1 and SphK2 in endothelial cells SphK1,2*^f/f^* and SphK1,2*^ECKO^* after 4-OHT (1μM, 72h) treatment (n=4/group). **(H)** SPT activity in SphK1,2*^f/f^* and SphK1,2*^ECKO^* after 4-OHT (1μM, 72h) treatment in absence or presence of S1P (300nM, 30’) or C16:0-ceramide (300nM, 30’) (n≥5/group from 3 independent EC isolations/group; 4 mice/EC isolation). **(I)** WB analysis of ORMDLs, NOGO-B, SPTLC1 and SPTLC2 in HUVEC lysates in absence or presence of S1P (300nM, 30’) and **(L)** relative ORMDLs quantification. (**M**) WB analysis of ORMDLs, SPTLC1 and SPTLC2 in HUVEC lysates in absence or presence of C16:0-ceramide (300nM, 30’), with or without SKI II 1µM, 1h pre-treatment). SPT activity and WB analysis of HUVEC treated with **(N)** siCTRL and siNOGO (40nM, 72h) (n≥4/group), or with **(O)** siCTRL and siORMDL1/2/3 (40nM, 72h) (n=4/group). **(P)** Graphical abstract. β-ACTIN, loading control. ^3^[H]-serine and palmitoyl-CoA were used as substrates for SPT activity. Sphinganine - the reaction product – was separated in TLC (thin-layer cromatography) and quantified. Where not indicated, data are representative of two or more independent experiments. Data are expressed as mean±SEM. \**P*≤0.05; \*\**P*≤0.01; \*\*\**P*≤0.001. Statistical significance was determined by unpaired t-test (**A-G, L**), and 2-way ANOVA with Tukey’s post-test (**H, N, O**).

### ORMDLs are degraded via PHD-mediated ubiquitination and proteasomal degradation

We next sought to unveil the molecular mechanism orchestrating the acute changes of ORMDLs levels. Protein abundance reflects the integration of synthesis and degradation rates^27^. The inhibition of translation with cycloheximide (CHX) showed that ORMDL half-life was 1.7h (**Fig. 2A**), indicating a relatively fast turnover. To identify the regulatory mechanisms controlling ORMDLs levels, we analyzed ORMDLs sequence properties and observed the presence in the C-terminus of a conserved prolyl hydroxylase (PHD) consensus domain, known to regulate the levels of hypoxia-inducible factor 1α (HIF-1α)^28^ and other proteins^29^ via ubiquitination (**Fig. 2B**). Interestingly, the inhibition of PHD activity with the hydroxylase inhibitor dimethyloxalylglycine (DMOG) elevated ORMDLs to the same extent as S1P, suggesting that PHD-mediated ubiquitination controlled the abundance of ORMDLs (**Fig. 2C**). A retro-translocation from the ER to the cytosol is necessary for ER-membrane associated protein degradation (ERAD)^30^. The inhibition of dislocase p97/VCP with Eeyarestatin-1 (EER1) led to ORMDLs accumulation (**Fig. 2D**), implicating the retro-translocation as regulatory step of ORMDLs levels. Eukaryotic cells rely on the ubiquitin-proteasome pathway as a major degradation system for short-lived proteins^31^. MG132, proteasome inhibitor, significantly augmented ORMDLs levels in basal conditions but not in presence of S1P, most likely because S1P already maximized ORMDLs stability by inhibiting their degradation (**Fig. 2E**). This data highlights proteasomal degradation as a primary mechanism to control mammalian ORMDLs levels.

**Figure 2.**
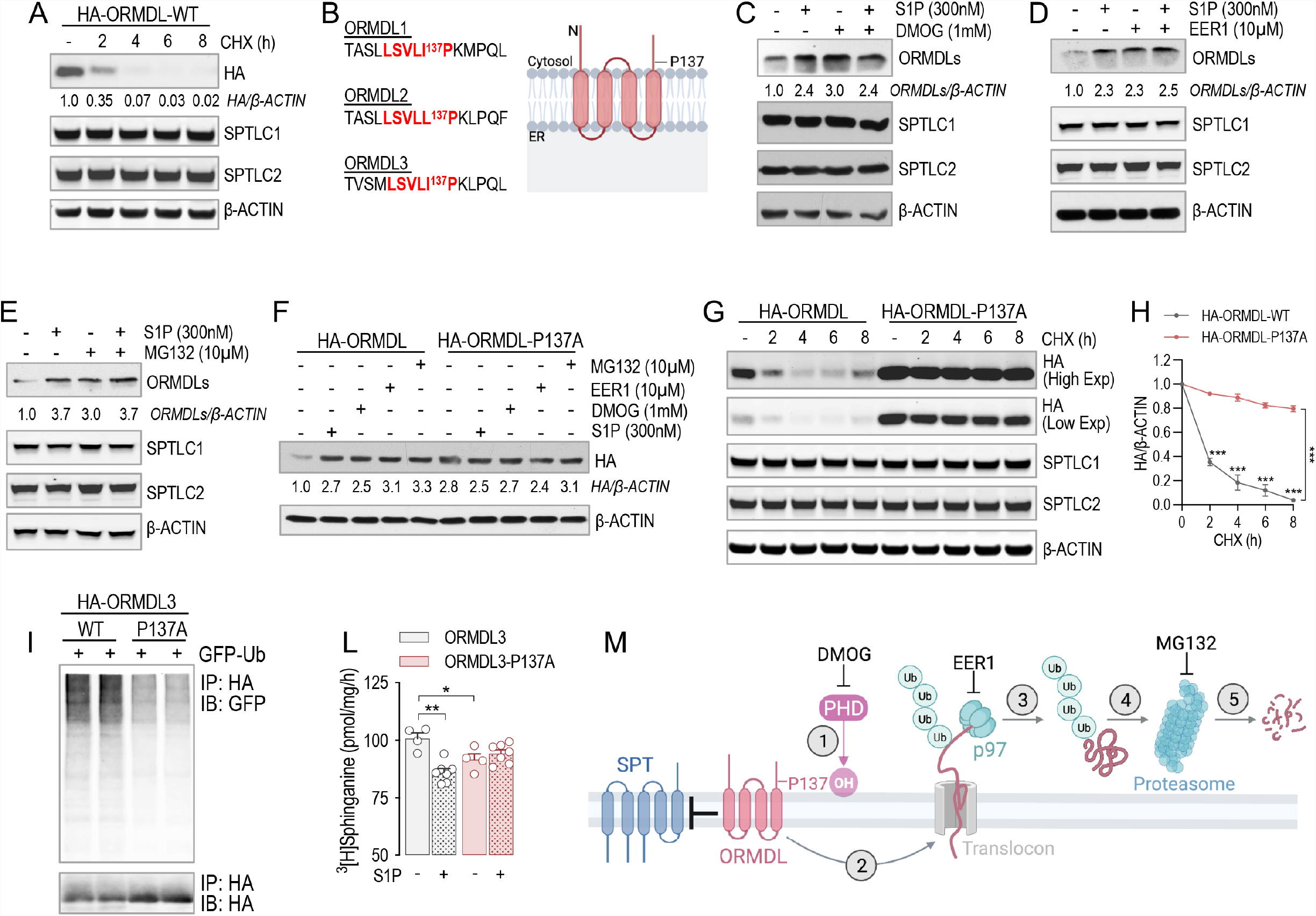
ORMDLs are degraded via PHD-mediated ubiquitination and proteasomal degradation. **(A)** WB analysis of ORMDL3 (HA), SPTLC1 and SPTLC2 in HUVEC lysates expressing HA-ORMDL3 and treated with cycloheximide (CHX, 10µM) for the indicated period of time. **(B)** Prolyl hydroxylase consensus sequence in the three ORMDL isoforms. **(C-E)** WB analysis of ORMDLs, SPTLC1 and SPTLC2 in HUVEC lysates in absence or presence of S1P (300nM, 30’) and with or without **(C)** DMOG (1mM, 1h pre-treatment), **(D)** Eeyarestatin 1 (EER1, 10µM, 1h pre-treatment), **(E)** MG132 (10µM, 1h pre-treatment). **(F)** WB analysis of HA-ORMDL3 and HA-ORMDL3-P137A in HUVEC lysates in absence or presence of S1P (300nM, 30’), DMOG (1mM, 1h), EER1 (10µM, 1h), and MG132 (10µM, 1h). **(G)** WB analysis of HA-ORMDL3 and HA-ORMDL3-P137A, SPTLC1 and SPTLC2 in HUVEC lysates treated with CHX (10µM) for the indicated period of time, and **(H)** relative quantification. **(I)** WB analysis for GFP and HA of HEK293T transfected with GFP-ubiquitin and with the indicated HA-ORMDL3 plasmid, and immunoprecipitated with HA antibody. **(L)** SPT activity in HUVEC expressing the ORMDL3 or ORMDL3-P137A, and depleted of endogenous ORMDLs with siRNA, in absence or presence of S1P (300nM, 30’) (n≥4/group). **(M)** Graphical abstract. β-ACTIN, loading control. ^3^[H]-serine and palmitoyl-CoA were used as substrates for SPT activity. Sphinganine - the reaction product – was separated in TLC (thin-layer cromatography) and quantified. Where not indicated, data are representative of two or more independent experiments. Data are expressed as mean±SEM. \**P*≤0.05; \*\**P*≤0.01; \*\*\**P*≤0.001. Statistical significance was determined by 2-way ANOVA with Tukey’s post-test.

ORMDL3 is the most abundant ORMDL isoform in EC (**Supplementary Fig. 3A**), and SNP for ORMDL3 are associated with asthma^6^ and atherosclerosis^7^. To demonstrate the requirement of PHD-mediated hydroxylation for ubiquitination-mediated degradation of ORMDLs, we mutated P137 to A in the PHD consensus domain of ORMDL3. The P137 to A mutation was sufficient to stabilize ORMDL3 to the same levels of S1P, DMOG, EER1, and MG132 (**Fig. 2F**). Interestingly, the half-life of ORMDL3-P137A was remarkably higher than the native form, 24h and 1.7h respectively (**Fig. 2G,H**). Of note, the expression of mutant ORMDL3 did not affect the stability of SPTLC1 or SPTLC2 (**Fig. 2G** and **Supplementary Fig. 3B,C**). Lys-48-linked ubiquitination targets protein for degradation^32^. WB analysis of immunoprecipitated ORMDL3 showed that Lys-48-linked polyubiquitination was significantly reduced in the ORMDL3-P137A compared to native ORMDL3 (**Fig. 2I**). Finally, to investigate the biological significance of P137 hydroxylation, native and mutant ORMDL3 were overexpressed in HUVEC depleted of endogenous ORMDLs via siRNA approach. In HUVEC expressing ORMDL3-P137A, SPT activity was significantly suppressed at baseline and no longer modulated by S1P (**Fig. 2L**). Mechanistically, these data strongly support the model in which ORMDLs undergo PHD-mediated hydroxylation of P137, ubiquitination, extraction from ER membrane via ERAD pathway, and ultimately proteasome-mediated degradation (**Fig. 2M**). S1P signaling downregulates SPT activity by stabilizing ORMDLs via inhibition of PHD-mediated hydroxylation.

### Endothelial-derived S1P inhibits de novo sphingolipid biosynthesis via S1PRs

Quiescent EC express mainly S1PR1, and less abundantly S1PR3^33^. The loss of either S1PR1 (**Fig 1A**) or S1PR3 (**Supplementary Fig. 4A**) leads to constitutive increases of SPT activity (**Fig. 1B** and **Supplementary Fig. 4B**), suggesting that S1PR1,3 signaling provides constitutive inhibitory feedback on SPT. Thus, to investigate the role of S1PR1,3 in the dynamic regulation of SPT activity by S1P, mEC were depleted of both S1P receptors. Interestingly, the loss of S1PR1,3 abolished the downregulation of SPT activity (**Fig. 3A**), as well as the stabilization of ORMDLs, by exogenous S1P (**Fig. 3B**), suggesting that these receptors are necessary to sense S1P abundance and operate a negative feedback to SPT via ORMDLs. Consistently, S1PR1,3 deletion significantly raised basal SPT activity (**Fig. 3A**), as well as total ceramides (**Fig. 3C**) and glucosylceramides (**Fig. 3D**) levels. Cellular SL results from the *de novo* and recycling pathways. To investigate the impact of S1P on SPT activity, we used stable-isotope labeled serine (L-serine-^13^C_3_,^15^N) to trace the *de novo* synthesized SL (**Fig. 3E**). Exogenous S1P significantly decreased labelled ceramides and glucosylceramides (**Fig. 3F-I**), corroborating the S1P-induced downregulation of SPT activity (**Fig. 1D**). However, in absence of S1PR1,3, labeled ceramides and glucosylceramides were significantly elevated in basal conditions and were not decreased by exogenous S1P compared to control EC (**Fig. 3F-I**), suggesting that S1PR1,3 are necessary to mediate S1P downregulation of SPT activity. Unlabeled ceramides showed a similar trend (**Supplementary Fig. 5A-E**), suggesting that the primary influence exerted on sphingolipid production was indeed from *de novo* sphingolipid biosynthesis.

**Figure 3.**
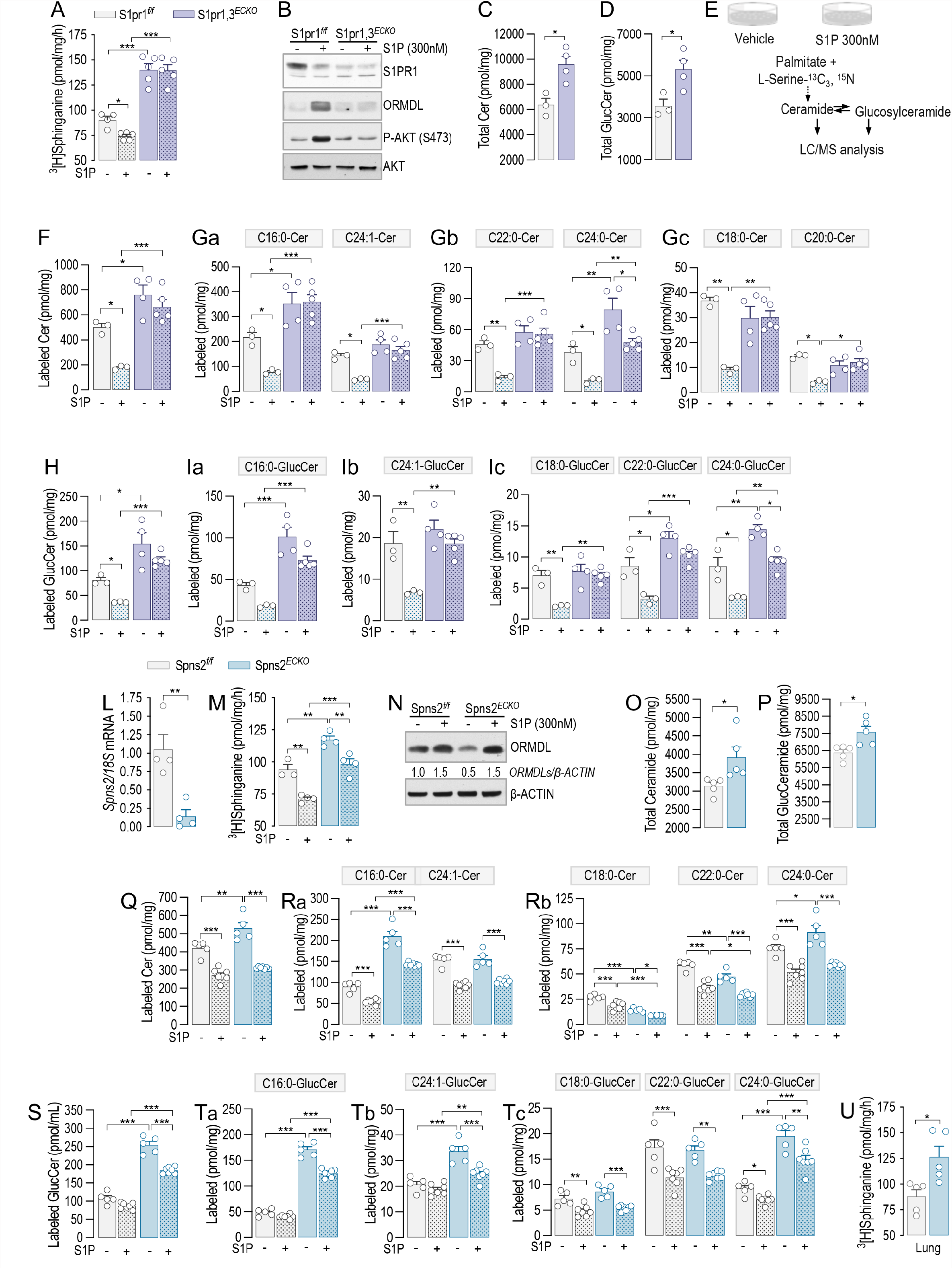
Endothelial-derived S1P inhibits de novo sphingolipid biosynthesis via S1PRs. **(A)** SPT activity in S1pr1,3*^f/f^* and S1pr1,2*^ECKO^* after 4-OHT (1μM, 72h) and siS1PR3 (40nM, 72h) treatments, in absence or presence of S1P (300nM, 30’) (n≥4/group from 2 independent EC isolations/group; 4 mice/EC isolation) and (**B**) relative WB analysis for S1PR1, ORMDL, P-AKT and AKT. LC-MS/MS quantification of Total **(C)** Ceramide and **(D)** Glucosylceramide in S1pr1,3*^f/f^* and S1Pr1,3*^ECKO^* endothelial cells after 4-OHT and siS1PR3 treatment (n≥3/group from 2 independent EC isolations/group; 4 mice/EC isolation). **(E)** Experimental procedure for the measurement of the *de novo* synthetized Ceramides and Glucosylceramides. LC-MS/MS quantification of **(F)** total and **(G)** specific Ceramides, and of **(H)** total and **(I)** specific Glucosylceramides labeled with L-Serine-^13^C_3_, ^15^N in absence or presence of S1P (300nM). **(L)** RT-PCR for Spns2 in endothelial cells Spns2*^f/f^* and Spns2*^ECKO^* after 4-OHT (1μM, 72h) treatment (n=4/group). **(M)** SPT activity in Spns2*^f/f^* and Spns2*^ECKO^* after 4-OHT, in absence or presence of S1P (300nM, 30’) (n≥3/group from 2 independent EC isolations/group; 4 mice/EC isolation) and (**N**) relative WB analysis for ORMDL. LC-MS/MS quantification of Total **(O)** Ceramide and **(P)** Glucosylceramide in Spns2*^f/f^* and Spns2*^ECKO^* endothelial cells after 4-OHT (n=5/group from 2 independent EC isolations/group; 4 mice/EC isolation). **(Q)** SPT activity in microsomes from lung of Spns2*^f/f^* and Spns2*^ECKO^* mice (n=5/group). LC-MS/MS quantification of **(R)** total and **(S)** specific Ceramides, and of **(T)** total and **(U)** specific Glucosylceramides labeled with L-Serine-^13^C_3_, ^15^N in absence or presence of S1P (300nM). β-ACTIN, loading control. ^3^[H]-serine and palmitoyl-CoA were used as substrates for SPT activity. Sphinganine - the reaction product – was separated in TLC (thin-layer cromatography) and quantified. Where not indicated, data are representative of two or more independent experiments. Data are expressed as mean±SEM. \**P*≤0.05; \*\**P*≤0.01; \*\*\**P*≤0.001. Statistical significance was determined by unpaired t-test (**C, D, L, O-Q**), and 2-way ANOVA with Tukey’s post-test (**A, F-I, M, R-U**).

Considering that the deletion of SphK1,2 dramatically upregulated SPT activity (**Fig. 1H**), we hypothesized that S1PR1,3 can be activated in an autocrine manner by locally produced S1P to initiate the negative feedback on SPT. To preserve the formation of intracellular S1P and its degradation by S1P lyase, representing the catabolic exit of the pathway, instead of *Sphk1,2* genes, *Spns2* was deleted to prevent the transporter-mediated cellular excretion of S1P (**Fig. 3L**). Thus, mEC from Spns2*^f/f^* VE-Cad-CreERT2 mice were used *in vitro*. Spns2 deletion significantly raised SPT activity, suggesting a constitutive inhibitory function of endogenous S1P on SPT via inside-outside signaling (**Fig. 3M**). Importantly, exogenous S1P was able to downregulate SPT activity in Spns2*^ECKO^* as in Spns2*^f/f^* EC (**Fig. 3M**), indicating that while the loss of Spns2 disrupted the autocrine S1P signal on SPT, the S1PR-ORMDL-SPT signaling pathway was preserved. Consistently with these findings, total ceramides (**Fig. 3O** and **Supplementary Fig. 5F,G**) and glucosylceramides (**Fig. 3P** and **Supplementary Fig. 5H-L**), were upregulated in Spns2*^ECKO^* vs. Spns2*^f/f^*. The measure of sphingolipid flux with isotope-labeled serine showed that the loss of Spns2 increased the *de novo* synthesized ceramides and glucosylceramides in basal conditions (**Fig. 3Q-T**), while preserving the downregulation in response to exogenous S1P (**Fig. 3Q-T**), in line with SPT activity data (**Fig. 3M**). Lastly, SPT activity was significantly higher in Spns2*^ECKO^* mouse lung microsome compared to Spns2*^f/f^* (**Fig. 3U**), corroborating the function of SPNS2-S1P negative feedback on SPT *in vivo*. These results support an important role of the endothelial-derived S1P-S1PR signaling in maintaining sphingolipid homeostasis via stabilization of ORMDL-SPT complex.

### Ceramide accrual leads to endothelial and mitochondrial disfunction

Ceramides regulate membrane biophysical properties, particularly of lipid rafts, important signaling platforms^34–36^. Recently, we reported that optimal ceramide levels are necessary to preserve endothelial signal transduction to different agonists, including VEGF and insulin^19^. The loss of Spns2 disrupted the cellular SL sensing mechanism and increased SL, including ceramides (**Fig 3O**). Thus, to test the hypothesis that in absence of Spns2 ceramide accrual impairs endothelial signal transduction, we performed a series of experiments in primary mEC *in vitro* isolated from Spns2*^ECKO^* and Spns2*^f/f^* mice. The activation of both VEGFR2 and insulin receptor (IR), and downstream signaling, were blunted in Spns2*^ECKO^* compared to Spns2*^f/f^* mEC (**Fig. 4A-C**), although the receptor expression was unchanged, suggesting that ceramide accrual in absence of Spns2 impairs endothelial signal transduction. To further explore the consequences of Spns2 deletion in more physiological setting, we used mesenteric arteries (MA) *ex vivo*. In line with endothelial signaling data, vasorelaxation to both VEGF (**Fig. 4D**) and insulin (**Fig. 4E**) was significantly diminished in Spns2*^ECKO^* compared Spns2*^f/f^* MA. On the contrary, acetylcholine and sodium nitroprusside (SNP) induced vasorelaxation were not affected by altered ceramide levels (**Fig. 4F,G**), as previously reported^19^. Interestingly, mouse treatment with myriocin, an inhibitor of SPT, restored VEGF- and Insulin-dependent vasorelaxation (**Fig. 4D,E**), consistent with a role of ceramide in endothelial disfunction^37^. Multiple lines of evidence have shown that ceramide accrual is causal of mitochondrial dysfunction, oxidative stress^38,39^, and apoptosis^40^. Loss of Spns2 resulted in decreased maximal respiration and reduced spare respiratory capacity (**Fig. 4H,I**). Extracellular acidification rate, an indirect index of mitochondrial dysfunction, suggests an increased ability to upregulate aerobic glycolysis upon the loss of mitochondrial ATP production caused by inhibition of the ATPase by oligomycin (**Fig. 4L,M**). Spns2*^ECKO^* EC showed decreased membrane potential (**Fig. 4N**), smaller mitochondrial size (**Fig. 4O**) and increased number of mitochondria per cell (**Fig. 4P**), as assessed by TMRM staining. All together, these findings suggests a respiratory chain dysfunction that result in lower mitochondrial polarization, fragmentation of mitochondria and adaptive changes in glycolysis as an alternate source of energy production, supporting the concept that Spns2 deletion increases sphingolipid de novo biosynthesis and ceramide levels, hence the susceptibility of the cells to metabolic stress.

**Figure 4.**
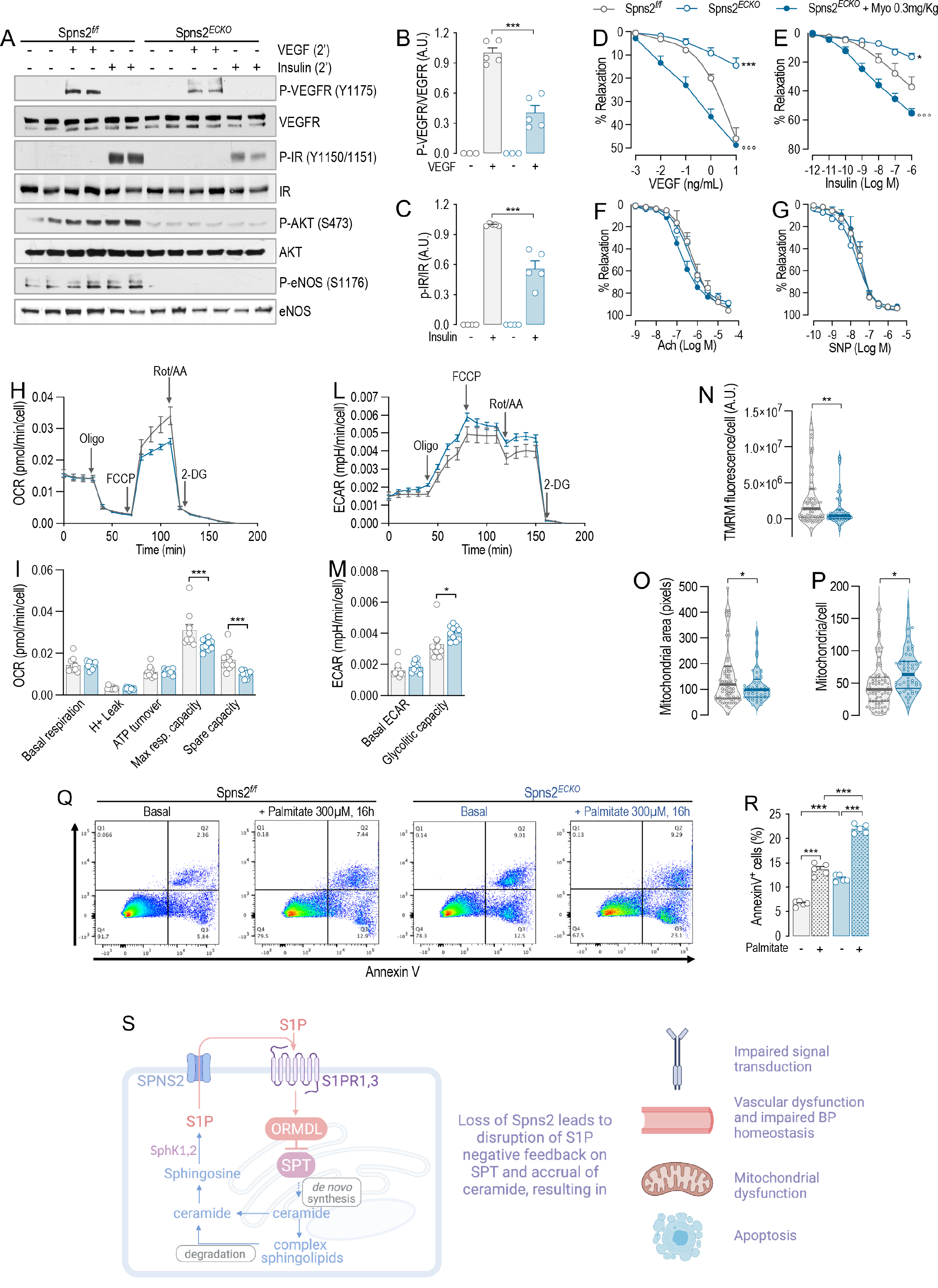
Ceramide accrual leads to endothelial and mitochondrial disfunction. **(A)** WB analysis of P-VEGFR2 (Y1175), VEGFR2, P-IR (Y1150/1151), IR, P-AKT (S473), AKT, P-eNOS (S1176), and eNOS in Spns2*^f/f^* and Spns2*^ECKO^* endothelial cells lysates in absence or presence of VEGF (100ng/mL, 2’) or Insulin (1U/mL, 2’) and **(B-C)** relative quantification of the indicated phospho/total protein ratios. Vasodilation in response to **(D)** VEGF, **(E)** Insulin, **(F)** Acetilcholine, and **(G)** SNP (Spns2*^f/f^*, n=3; Spns2^ECKO^, n=3; Spns2^ECKO^ + Myo, n=4). **(H)** Representative curves of OCR and **(I)** quantification of OCR metrics in Spns2*^f/f^* and Spns2*^ECKO^* mEC. Oligomycin (Oligo), uncoupler FCCP, rotenone and antimycin A (Rot/AA) were added at the indicated times. **(L)** Representative curves of ECAR and **(M)** quantification of ECAR metrics in Spns2*^f/f^* and Spns2*^ECKO^* mEC. Oligomycin (Oligo), uncoupler FCCP, rotenone and antimycin A (Rot/AA) were added at the indicated times. **(N)** TMRM fluorescence, (**O**) mitochondrial area, and (**P**) number of mitochondria per cells, as quantified by TMRM and Hoechst fluorescence. **(Q)** Representative dot-plot diagrams and **(R)** relative quantification of Spns2*^f/f^* and Spns2*^ECKO^* mEC, in absence or presence of Palmitate (300µM, 16h), stained with Annexin V and analyzed by FACS. Where not indicated, data are representative of two or more independent experiments. Data are expressed as mean±SEM. \**P*≤0.05; \*\**P*≤0.01; \*\*\**P*≤0.001; °°°*P*≤0.001. *Spns2*^f/f^* vs. Spns2*^ECKO^*, °Spns2*^ECKO^* vs. Spns2*^ECKO^* + Myo. Statistical significance was determined by 2-way ANOVA with Tukey’s post-test (**D-G, I, M, R**) and unpaired t-test (**N, O, P**).

Next, we investigated whether the ceramide accrual in absence of Spns2 heighten the susceptibility of the EC to apoptosis induced by palmitate, substrate of SPT. In Spns2*^ECKO^* EC baseline apoptosis was increased to the same extent of palmitate-induced apoptosis in Spns2*^f/f^* EC. The loss of Spns2 dramatically increased the apoptosis induced by palmitate (**Fig. 4P,Q**).

Altogether, these results show that disruption of S1P-S1PR-ORMDL negative feedback on SPT results in unrestricted levels of SL, including ceramides, triggering mitochondrial dysfunction, apoptosis, and impaired vascular tone regulation, which all underline endothelial dysfunction, an early event in the pathogenesis of cardiovascular diseases, including atherosclerosis.

## DISCUSSION

SL, a minor class of mammalian lipids, have gained much attention because emerging pre-clinical and clinical evidence established a strong link between altered sphingolipid homeostasis and diseases^41,42^, including atherosclerosis^43^, coronary artery disease^44^, myocardial infarction^45^, heart failure^46,47^, hypertension^48^, and type 2 diabetes^49,50^. However, how cells sense sphingolipid levels and regulate their biosynthesis accordingly remains poorly understood. Because SPT catalyzes the rate-limiting step of the pathway, there has been much effort in understanding how this enzyme is modulated in response to cellular sphingolipid levels. In this study, we identified a sensing mechanism used by the cells to maintain sphingolipid homeostasis and assure proper cellular functions. Our study discovered S1P as the specific SL metabolite sensed by the cells; S1P actions downregulate SPT activity via S1PR/ORMDL negative feedback to maintain sphingolipid homeostasis.

In yeast the phosphorylation of Orms releases the interaction with SPT, hence sphingolipid *de novo* biosynthesis is upregulated^2^. However, in mammalian orthologs ORMDLs lack the N-terminus hosting the phosphosites^9^. Considering that sphingolipid homeostasis is necessary to preserve cellular functions and health, it is conceivable that complex regulatory mechanisms exist to regulate SPT-ORMDL interactions in response to metabolic and environmental cues, at transcriptional, translational and post-translational levels. For instance, inflammatory stimuli and changes in SL can upregulate the transcription^51^ and the translation^25^ of ORMDL, respectively, although the underlying molecular mechanisms remain unknown. Previous works from Wattenberg’s group^24,26^, using cell-free isolated membranes or permeabilized cells, suggested that C6:0- and C8:0-ceramide (10-20μM) can acutely downregulate SPT activity, probably by interacting with the SPT-ORMDLs complex. However, by using multiple genetic and pharmacological strategies our data showed that when S1P formation was inhibited, ceramide was no longer able to modulate SPT activity.

ORMDLs have a relatively short half-life (ca. 1.7h, **Fig. 2A,H**). Mechanistically, our data show that PHD-mediated hydroxylation of Pro137 enforces a constitutive degradation of ORMDLs via ER-associated degradation and ubiquitin-proteasome pathway (**Fig. 2M**), hence preserving a steady-state SPT activity. Upregulation of S1P signaling inhibits PHD-mediated Pro137 hydroxylation, resulting in ORMDLs stability and downregulation of SPT activity. It is conceivable that this sensing-effector mechanism that maintains sphingolipid homeostasis is not only functional in the EC but also in other cell types. Further studies are needed to explore this possibility.

S1P is known to induce barrier function^18^, NO production^3,14,15^, as well as cell migration and survival. In addition to known functions, our study discovered that a fundamental function of S1P is to maintain sphingolipid homeostasis via S1PR-ORMDL negative feedback on SPT. Genetic and pharmacological disruption of endothelial S1P autocrine signaling at multiple levels, including *Sphks*, *Spns2*, and *S1prs*, support the direct role of S1P-S1PR signaling in modulating SPT activity via stabilization of ORMDLs. How cells can sense SL has been a longstanding question. Our findings identified S1P-S1PR as the sensor-effector unit by which cells can sense SL and maintain homeostasis by a nimble modulation of SPT activity via ORMDLs.

The disruption of S1P negative feedback on SPT leads to uncontrolled sphingolipid *de novo* biosynthesis and ceramide accumulation, resulting in mitochondrial dysfunction, apoptosis, and impaired signal transduction and endothelial-regulated vascular tone, which are all manifestations of endothelial dysfunction, an early event in the onset of cardiovascular diseases, including atherosclerosis and hypertension. Postnatally the disruption of S1P signaling results in permeability^18^, hypertension^15,23^, atherosclerosis^16^, and heart failure^52^, as result of the loss of S1P biological actions, including the enhancement of endothelial barrier functions and NO production, and downregulation of NFkB pathway^16^. In addition to these canonical cardiovascular functions, our data identified a novel fundamental function of S1P which is to maintain sphingolipid homoeostasis and protect the cells from a “metabolic catastrophe” due to uncontrolled ceramide accrual, causing cellular and organ dysfunctions.

The clinical significance of this signaling mechanisms is also underlined by the correlation of single nucleotide polymorphisms (SNP) in *SphK1*^53^, *S1pr1*^54^, and *Ormdl3*^7^ with the risk of atherosclerosis. Genetic deletion of endothelial *S1pr1* in mice significantly elevated BP^15^. Interestingly, chronic administration fingolimod, a functional antagonist of S1PR1 approved by the FDA for the treatment of relapsing remitting multiple sclerosis^55^, also significantly increased BP, in line with the loss endothelial *S1pr1*^15^. Our findings suggest that an additional biological consequence of fingolimod on-target actions on S1PRs is the disruption of S1P negative signaling on SPT, and therefore the homeostasis of SL, at least in cell type expressing high levels of S1PR1, such as endothelial and immune cells^13,33^. Further studies are needed to investigate the impact of fingolimod on sphingolipid sensing and homeostatic pathway of the cells.

This study reveal a novel mechanism by which cells can sense SL and regulate sphingolipid biosynthesis accordingly. S1P is the metabolite sensed by the cells to downregulate SPT activity via S1PR-ORMDl signaling. Whether the disruption of this negative feedback plays a role in the pathogenesis human diseases remains to be investigated.

## MATHERIAL AND METHODS

### Mouse models

We generated conditional mouse model lacking endothelial S1PR1, namely S1pr1*^ECKO^*; lacking endothelial SphK1,2, namely SphK1,2*^ECKO^*; and lacking endothelial Spns2, namely Spns2*^ECKO^*. S1pr1*^f/f^* (floxed S1pr1)^56^, SphK1,2*^f/f^* (floxed SphK1,2)^57^, and Spns2*^f/f^* (floxed Spns2)^58^ mice were crossed with transgenic mice in which the VE-cadherin promoter drives expression of tamoxifen-responsive Cre (VE-Cad-CreERT2), such that tamoxifen treatment selectively deletes the *lox*P-flanked (floxed) region of S1pr1 in endothelial cells (EC)^59^. S1pr1*^ECKO^* and S1pr1*^f/f^*, and SphK1,2*^ECKO^* and SphK1,2*^f/f^* were used only for isolation of mouse liver EC. To delete Sptlc2 in EC, 7- to 8-week-old male mice were injected intraperitoneally with 20 mg/kg of tamoxifen daily for 5 consecutive days. All animal experiments were approved by the Weill Cornell Institutional Animal Care and Use Committee.

### Isolation of mouse liver EC

Liver EC, but not lung and heart EC, are responsive to 4-hydroxytamoxyfen-induced gene excision *in vitro*. Four-week-old female and male mice were used to isolate EC. Briefly, liver were cut into small pieces and incubated in a solution of 2mg/mL collagenase I (Alfa Aesar, #J62406), 1U/mL dispase (Stemcell Technologies, #07913) and 100µg/mL DNase I (Roche, # 10104159001), followed by mechanical dissociation. EC were isolated with CD144 antibody-conjugated dynabeads (CD144 antibody, BD bioscience, #555289; dynabeads, ThemoFisher Scientific, #11035). Isolated EC were cultured in DMEM (Lonza, #12709-F) with 20% FBS (R&D Systems, #S11150H), 100μg/mL heparin (Sigma, #H3393), and 25μg/mL ECGF (Alfa Aesar, #J64516). For gene excision, EC were treated with 1μM 4-hydroxytamoxyfen (Cayman Chemical, #14854) for 3 consecutive days. Before treatment with S1P (300nM, 30min; Cayman Chemical, #62570), C16:0-ceramide (300nM, 30min; Avanti, #868516), VEGF (100ng/mL, 2min; Peprotech, #100-20), or Insulin (1U/mL, 2min), ECs were cultured in DMEM with 10% Charcoal-Stripped FBS for 18h, followed by 6h starvation in DMEM with 0.1% Charcoal-Stripped FBS.

### Experimental protocol with HUVEC

HUVEC (LifeLine Cell Technology, cat# FC-0044) were grown in EBM-2 (Lonza, cat# CC-3156) and supplemented with EGM-2 Endothelial Cell Growth Medium-2 BulletKit (Lonza, cat# CC-3162) and 10% FBS. Before treatment HUVEC were cultured in EBM-2 with 10% Charcoal-Stripped FBS for 18h, followed by 6h starvation in EMB-2 with 0.1% Charcoal-Stripped FBS. The following treatments were used: S1P (300nM, 30min), C16:0-Ceramide (300nM, 30min), SKI II^60^ (1μM, 30min before C16:0-Cer; Cayman Chemical, #10009222), CHX (10μM, for the indicated time; Cayman Chemical, #14126), DMOG^28^ (1mM, 1h; Cayman Chemical, #72210), Eeyarestatin 1^61^ (EER1, 10μM, 1h; Cayman Chemical, #10012609); MG132 (10μM, 1h; Cayman Chemical, #13697).

### SPT activity assay

SPT activity in and HUVEC and EC was measured as previously described^19^. Briefly, the assay was conducted in 0.1mL of SPT reaction buffer composed by: 0.1M HEPES pH 8.3, 5mM DTT, 2.5mM EDTA, 50μM pyridoxal 5′-phosphate (Sigma, #P9255), 0.45μM [^3^H]serine (American Radiolabeled Chemicals, #0246), 0.2mM palmitoyl-CoA (Sigma, #P9715) and 150μg of protein lysates. After 15 min at 37 °C, the reaction was stopped with NH_4_OH and the reaction product 3-ketosphinganine converted into sphinganine with NaBH_4_ (5 mg/ml). Radiolabeled lipids were extracted by using a modified Bligh and Dyer’s method, dissolved in CHCl_3_ and analyzed by thin-layer chromatography.

### Western Blot analysis

RIPA buffer cell lysates were analyzed with sodium dodecyl sulfate–polyacrylamide gel electrophoresis (SDS-PAGE) and immunoblotting, as previously reported ^19^. The following primary antibodies were used for WB analysis: S1P1 (ABclonal, #A12935); ORMDL3 (Millipore, #ABN417); NOGO-B (SCBT, #sc-11027); SPTCL1 and eNOS (BD Biosciences, #611305 and (#610297, respectively); SPTLC2 (ABclonal, #A11716); HA, Ubiquitin-K48, P-IR (Y1150/1151), IR, P-VEGFR2 (Y1175), VEGFR2, P-eNOS (S1177), P-AKT (S473), and AKT (Cell Signaling Technology, #3724, #4289, #3024, #3020, #2478, #2479, #9571, #4058, and #2920, respectively); β-ACTIN (ThermoFisher Scientific, #AM4302).

### Phosphorylation analysis by phosphate-affinity SDS–PAGE

RIPA buffer EDTA-free cell lysates were loaded in a SDS–PAGE gels with prepared with 50mM MnCl_2_ and 25mM phosphate affinity reagent (ApexBio, #F4002)^62^. Gels were run at 100V for 2h, rinsed three time for 10min in transfer buffer with 1mM EDTA before transfer to nitrocellulose membranes.

### Knockdown by siRNA transfection

siRNA targeting Nogo-B (sense: 5’-GACUGGAGUGGUGUUUGGUUU-3’, antisense: 5’-ACCAAACACCACUCCAGUCUU-3’); ORMDL1 (sense: 5’-CUCAUUGGGAACAACUGGAUU-3’, antisense: 5’-UCCAGUUGUUCCCAAUGAGUU-3’); ORMDL2 (sense: 5’-CUUCCUUCAUACGGUGAA AUU-3’, antisense: 5’-UUUCACCGUAUGAAGGAAGUU-3’); ORMDL3 (sense: 5’-UUCUACACUAAGUACGACCUU-3’, antisense: 5’-GGUCGUACUUAGUGUAGAAUU-3’); S1P3 (sense: 5’-GCUCCAGUAACAACAGCAGUU-3’, antisense: 5’-CUGCUGUUGUUACUGGAGCUU-3’), and control (sense: 5’-UUCUCCGAACGUGUCACG-3’, antisense: 5’-ACGUGACACGUUCGGAGAA-3’) were synthesized by Dharmacon. HUVEC or murine EC were transfected with 40nM of siRNA using DharmaFECT 4 transfection reagent (Dharmacon, #T-2004). mRNA or protein expression and relative assays were performed 72h after transfection.

### Real-time PCR (RT-PCR) analysis of murine EC

Total RNA from EC in culture was extracted according to the TRIzol reagent protocol (ThermoFisher Scientific, #15596026). Maxima First Strand cDNA Synthesis Kit (ThermoFisher Scientific, #K1641) was used for the reverse transcription of 100ng of RNA. For RT-PCR analysis PowerUp™ SYBR™ Green Master Mix (ThemoFisher Scientific, #A25779) and Applied Biosystems 7500 Fast RT PCR system were used. Primers set were: SphK1 (5’-AGGTGGTGAATGGGCTAATG-3’ and 5’-TGCTCGTACCCAGCATAGTG-3’); SphK2 (5′-TGGTGCCAATGATCTCTGAA-3′ and 5′-CCAGACACAGTGACAATGCC-3′); ORMDL1 (5’-CATAGCCGGTTGAAGCAGAC-3’ and 5’-ACGTTGACTCAGAGCCTTGA-3’): ORMDL2 (5’-CCAAGTACGATGCTGCTCAC-3’ and 5’-TTCCAGTGCCTTCCCTCAAT-3’); ORMDL3 (5’-ACTGAGGTTGTAGCCCCTTC-3’ and 5’-ACCCTAACCCCACTACAAGC-3’); S1PR3 (5’-GCTTCATCGTCTTGGAGAACCTG-3’ and 5’-CAGAGAGCCAAGTTGCCGATGA-3’); Spns2 (5’-AGAAGCCGCATCCTCAGTTAGC-3’ and 5’-CAGGCCAGAATCTCCCCAAATC-3’); 18S (5’-TTCCGATAACGAACGAGACTCT-3’ and 5’-TGGCTGAACGCCACTTGTC-3’). Gene of interest relative mRNA expression was calculated with the 2(−ΔΔCt) method, using 18S as housekeeping^63^.

### Lentivirus construction

Human HA-tagged ORMDL3 (NCBI AAM43507.1) and the mutant P137A were synthesized by Genewiz and inserted in the lentiviral vector pCDH-CMV-MCS-EF1-Puro (Addgene). Lentiviral particles containing the construct encoding HA-ORMDL3-WT and HA-ORMDL3-P137A were produced in HEK293T cells transfected with Calcium Phosphate technique. Viral particles were harvested from the culture supernatant 72h after transfection, passed through a 0.45μm filter and concentrated by adding a virus precipitation solution (40% PEG8000 and 2.5M NaCl) overnight at 4 °C, followed by centrifugation at 1,500×*g* for 30min. Viral pellets were resuspended in DMEM and stored at −80 °C until use.

### Immunoprecipitation

To asses ubiquitination, 293T cells were co-transfected with plasmid expressing Ub-GFP (gift from Nico Dantuma (Addgene plasmid # 11928; http://n2t.net/addgene:11928; RRID:Addgene_11928)) and HA-ORMDL3-WT or HA-ORMDL3-P137A. After 48h, cells were lysed and HA-ORMDL3 was immunoprecipitated with antibody against HA (Cell Signaling Technology, #3724) in modified RIPA buffer (50 mM Tris-HCl pH 7.2, 0.9% NaCl, 5.0 mM NaF, 1.0 mM Na_3_VO4, 1% NP40, and protease inhibitors) at 4°C o.n. The immune complexes were precipitated with Dynabeads protein G (#10003D, Invitrogen, for 1.5h at 4°C) and size-fractionated on SDS-PAGE gels. Ubiquitin was detected with Ubiquitin-K48 antibody (Cell Signaling Technology, #4289).

### Microsomal isolation from mouse lung

Microsomal fractions were obtained from Spns2*^f/f^* and Spns2*^ECKO^* lungs as previously described ^3^. Briefly, lungs were homogenized with liquid nitrogen in microsomal preparation buffer (50mM HEPES pH 7.4, 0.25M sucrose, and 5mM EDTA). The homogenates were centrifuged for 15min at 18,000×*g* at 4°C, and the resulting supernatants were ultracentrifuged for 1h at 100,000×*g* at 4°C. The microsomal pellets were then resuspended by adding 0.25ml of SPT reaction buffer.

### Measurement of Sphingolipid Flux using Stable Isotopes

Confluent EC were switched to DMEM lacking L-Serine for 2hrs. The cells were then switched to DMEM containing 0.45mM L-serine-^13^C_3_,^15^N (Sigma, #608130) and 300μM palmitate, with or without 300nM S1P for 3h. The reaction was thereafter washed with PBS, trypsinized and the cell pellet stored at −80°C. Lipids were thereafter extracted and total and labeled sphingolipids analyzed by mass-spectrometry at the University of Utah metabolomics core, as previously described^64^.

### Flow cytometric determination of apoptosis by Annexin V/Propidium Iodide staining

Cells were analyzed for phosphatidylserine exposure by an Annexin-V FITC/Propidium Iodide double-staining method, according to manufacturer instruction (Abcam, #ab14085). Spns2*^f/f^* and Spns2*^ECKO^* cells were treated with vehicle or C6:0-ceramide (30μM, 8h; Avanti, #860506), stained, acquired with BD FACSymphony Flow Cytometer, and analyzed with FlowJo software.

### Vascular reactivity studies

At 2 weeks post-tamoxifen treatment, second order mesenteric arteries (MA) were harvested, cleaned from adhering tissue and mounted on glass micropipettes in a wire myograph chamber (Danish MyoTechnology, Aarhus, Denmark). Vessels were maintained in Krebs solution^3,19^. MA were equilibrated for 15 min at 80 mmHg, pre-constricted with PE (1μM) and a cumulative concentration-response curve of Ach (0.1nM-30μM) was performed to evaluate the endothelial function. The following concentration-response curves were performed: insulin (pU/mL-3μU/mL), VEGF (1μg/ml-30mg/ml), and sodium nitroprussiate (SNP, 10nM-30μM). Were indicated, mice were treated with myriocin 0.3mg/Kg i.p. for two consecutive days before the experiment.

### Seahorse

Oxygen consumption rate (OCR) and extracellular acidification rate (ECAR) was measured with a XF96 Extracellular Flux Analyzer (Agilent Technologies, Santa Clara, CA, USA). mEC were plated a density of 1.5 × 10^4^ cells/well in 200μl of DMEM and incubated for 24h at 37°C in 5% CO_2_. After replacing the growth medium with 200μl of XF Assay Medium (Seahorse Bioscience, 103575-100) supplemented with 5mM glucose, 1mM pyruvate and 2mM GlutaMAX (Gibco), pre-warmed at 37°C, cells were preincubated for 1h before starting the assay procedure. OCR and ECAR were recorded at baseline, in the presence of 1µM oligomycin, 2µM carbonyl cyanide 4-trifluoromethoxyphenylhydrazone (FCCP), 0.5µM Antimycin A (AA) plus 0.5µM Rotenon (Rot) and in the presence of 25mM 2-Deoxy-D-glucose sequentially. Non-mitochondrial respiration (in the presence of AA+Rot) was subtracted from all rates. Following the experiment cell nuclei were stained with 1μM Hoechst 33342 (Thermo), imaged with an ImageXpress pico (Molecular Devices, San Jose, CA, USA) and counted. OCR and ECAR were normalized by the cell counts. Respiratory and glycolysis parameters were quantified by subtracting respiration rates at times before and after the addition of electron transport chain inhibitors according to Seahorse Bioscience. Basal respiration: baseline respiration minus (AA+Rot)-dependent respiration; H+ leak, Oligo-dependent respiration minus (AA+Rot)-dependent respiration; ATP turnover, baseline respiration minus oligo-dependent respiration; Max respiratory capacity: FFCP–dependent respiration minus (AA+Rot)-dependent respiration; Spare capacity: Max respiratory capacity minus Basal respiration. Basal glycolysis: basal ECAR minus non-glycolitic acidification; Glycolitic capacity: maximal ECAR after oligo minus non-glycolitic acidification.

### Mitochondrial membrane potential and morphology

For the measurements of mitochondrial membrane potential number, mEC were seeded at the density of 2 × 10^3^ cells/well in a 96-well glass bottom tissue culture plate (Cellvis P96-1.5H-N) in 200μL of DMEM and incubated for 24h at 37°C in 5% CO_2_. Cells were loaded with 15nM tetramethylrhodamine methyl ester (TMRM, 544ex; 590em, Life Technologies) and 1μM Hoechst 33342 for 30 minutes at 37°C in Krebs buffer. TMRM and Hoechst fluorescence were imaged with an ImageXpress pico. Subsequently 5μM FCCP was added to record background fluorescence. ∼50 cells were segmented manually using ImageJ. Background TMRM fluorescence was subtracted from baseline fluorescence. Mitochondria were segmented and counted in each cell using adaptive thresholding.

## Statistical Analysis

Two-way ANOVA with Tukey’s post-test, or Student *t* test were used for the statistical analysis as indicated in figure legends. Differences were considered statistically significant when *P*<0.05. GraphPad Prism software (version 9.0, GraphPad Software, San Diego, CA) was used for all statistical analysis.

## DATA SOURCE

Figure 1 – Source Data 1: Uncropped western blot images

Figure 1 – Source Data 2: Ceramide and Glucosylceramide measurement in S1pr1*^f/f^* and S1pr1*^ECKO^* endothelial cells

Figure 2 – Source Data 1: Uncropped western blot images

Figure 3 – Source Data 1: Uncropped western blot images

Figure 3 – Source Data 2: Ceramide and Glucosylceramide measurement in S1pr1,3*^f/f^* and S1pr1,3*^ECKO^* endothelial cells

Figure 3 – Source Data 3: Ceramide and Glucosylceramide measurement in Spns2*^f/f^* and Spns2*^ECKO^* endothelial cells

Figure 4 – Source Data 1: Uncropped western blot images

Supplementary Figure 4 – Source Data 1: Ceramide and Glucosylceramide measurement in S1pr1,3*^f/f^* and S1pr3*^ECKO^* endothelial cells

Supplementary Figure 5 – Source Data 1: Ceramide and Glucosylceramide measurement in S1pr1,3*^f/f^* and S1pr1,3*^ECKO^* endothelial cells

Supplementary Figure 5 – Source Data 2: Ceramide and Glucosylceramide measurement in Spns2*^f/f^* and Spns2*^ECKO^* endothelial cells

---

**Supplementary Figure 1.**
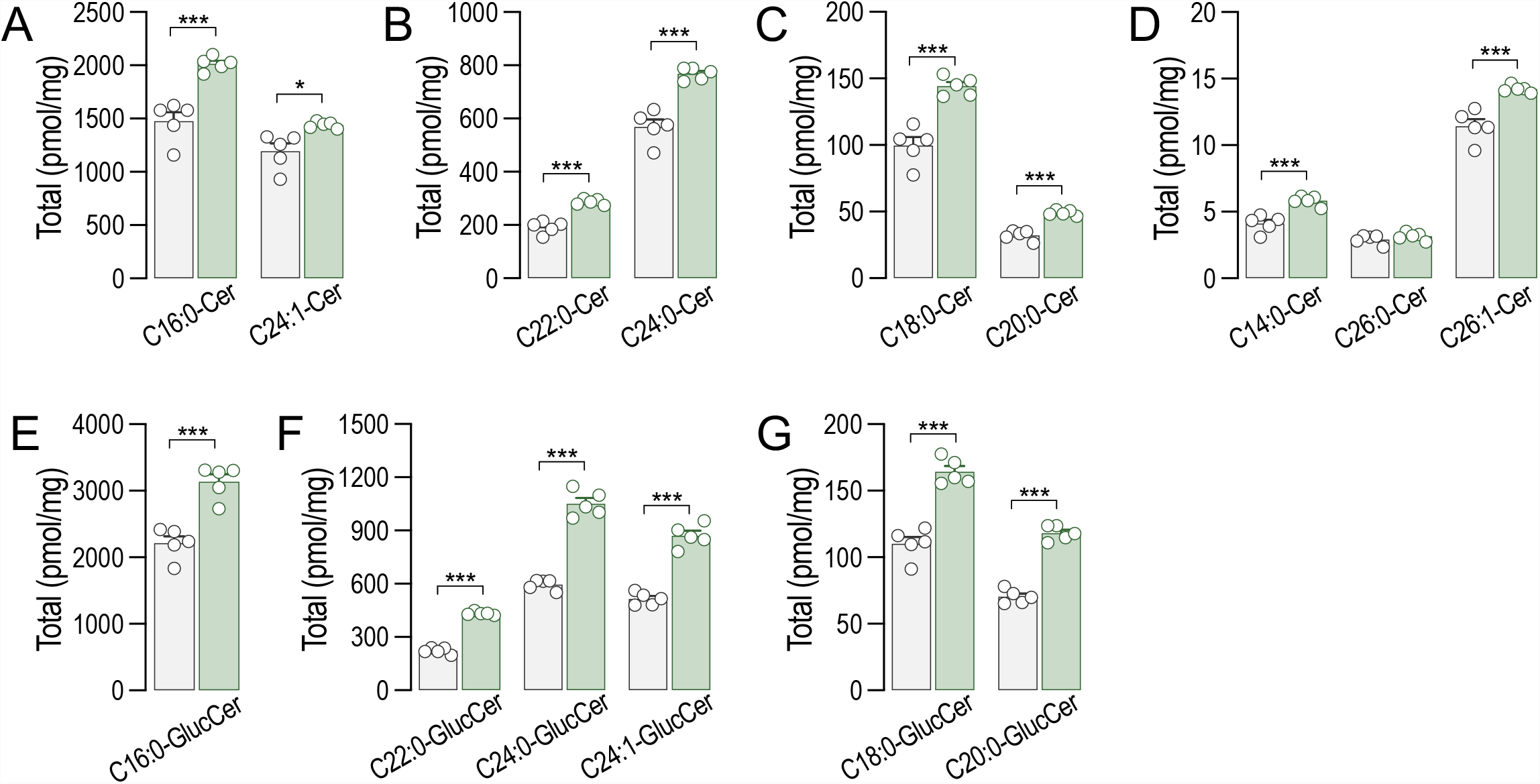
LC-MS/MS quantification of total specific **(A-D)** ceramides and **(E-G)** glucosylceramides in S1pr1*^f/f^* and S1Pr1*^ECKO^* endothelial cells after 4-OHT (1μM, 72h) treatment (n=5/group from 2 independent EC isolations/group; 4 mice/EC isolation). Data are expressed as mean±SEM. \*\*\**P*≤0.001. Statistical significance was determined by unpaired t-test.

**Supplementary Figure 2.**
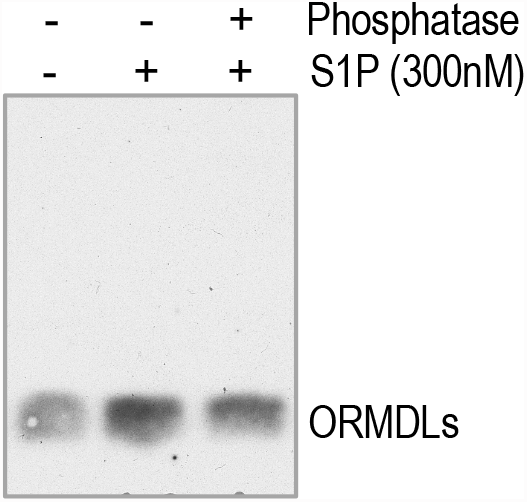
Phosphate-affinity gel analysis of ORMDLs in HUVEC in absence or presence of S1P (300nM, 30’).

**Supplementary Figure 3.**
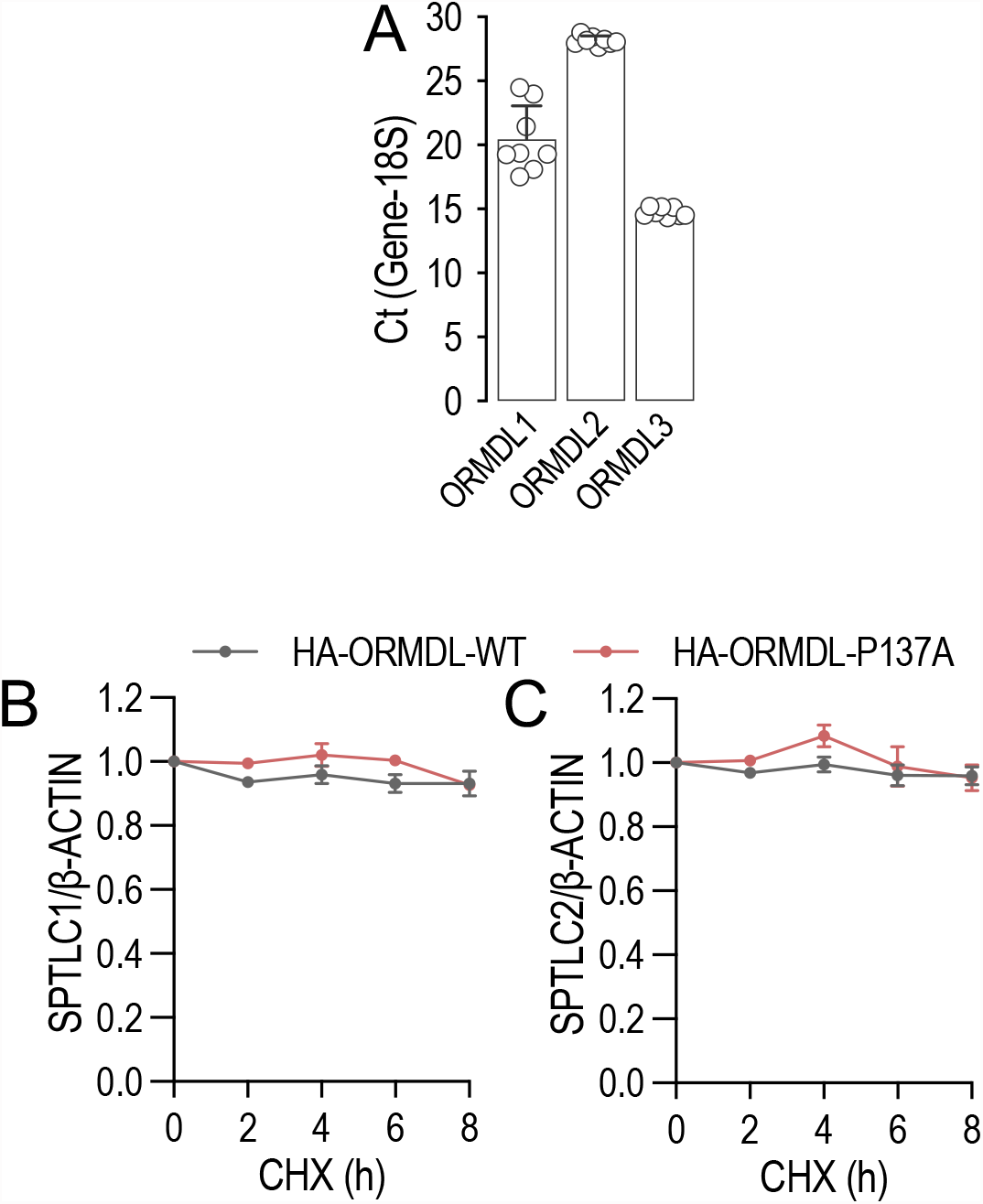
**(A)** Ct differences between genes and 18S determined by RT-PCR. Quantification of **(A)** SPTLC1, and **(B)** SPTLC2 (relative to Fig. 2G) in HUVEC lysates treated with CHX (10μM) for the indicated period of time, expressing HA-ORMDL3 or HA-ORMDL3-P137A. Data are expressed as mean±SEM. Statistical significance was determined by 2-way ANOVA with Tukey’s post-test.

**Supplementary Figure 4.**
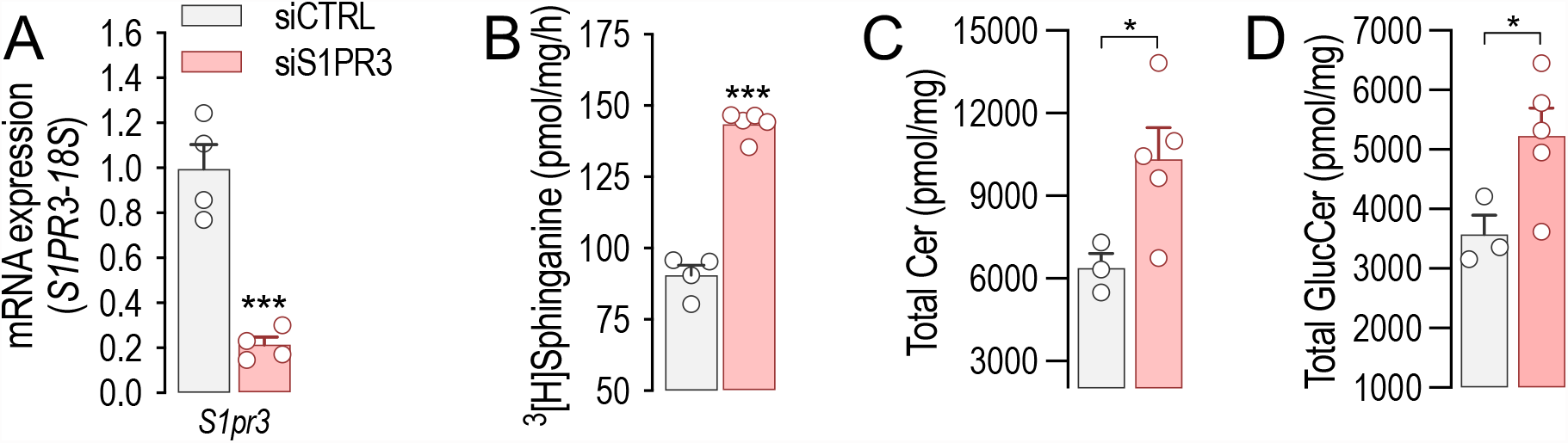
**(A)** RT-PCR and **(B)** SPT activity for S1pr3 in mEC after siCTRL or siS1PR3 treatment (40nM, 72h) (n=4/group). LC-MS/MS quantification of total **(C)** Ceramide and **(D)** Glucosylceramide in mEC after siCTRL or siS1PR3 treatment (40nM, 72h) (n≥3/group). Data are expressed as mean±SEM. \**P*≤0.05; \*\*\**P*≤0.001. Statistical significance was determined by unpaired t-test.

**Supplementary Figure 5.**
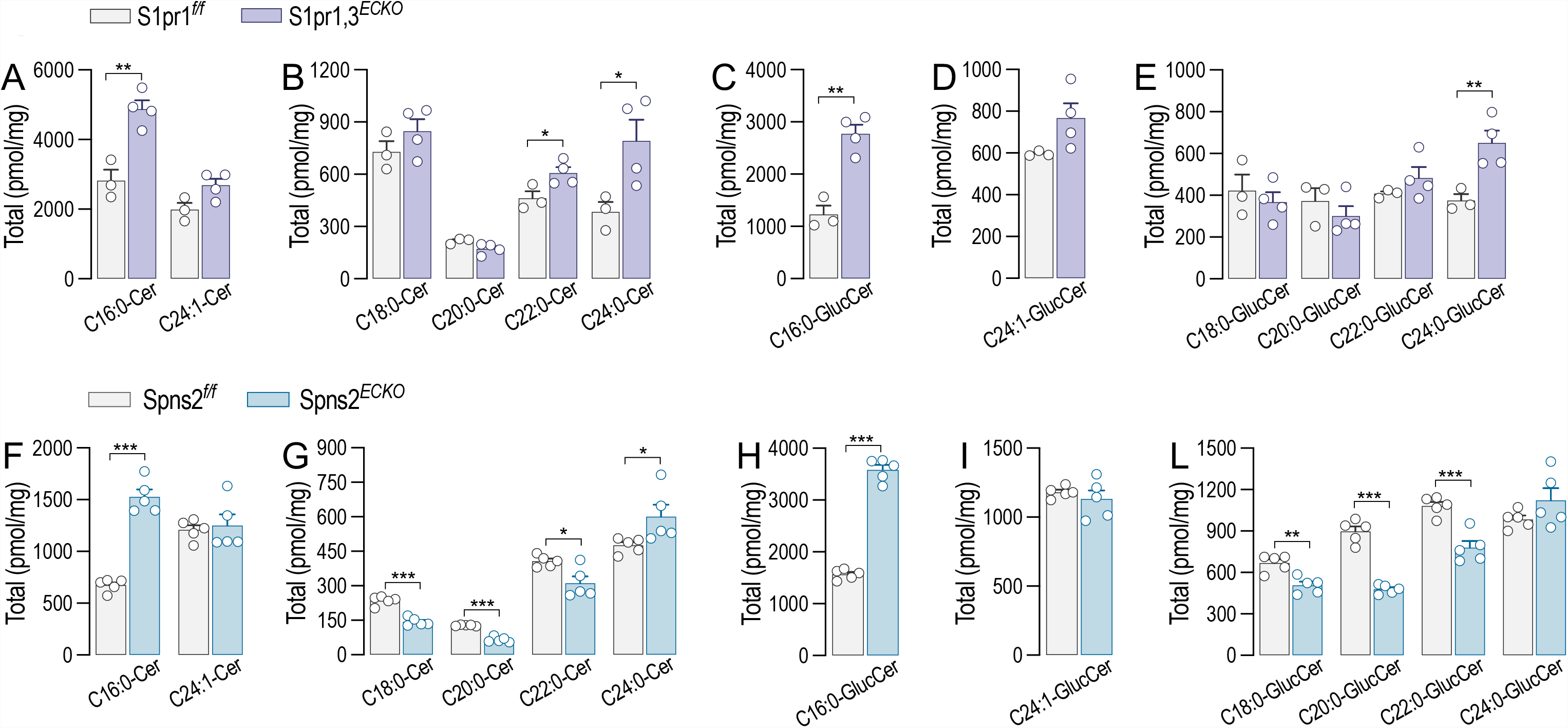
LC-MS/MS quantification of total specific **(A-B)** ceramides and **(C-E)** glucosylceramides in S1pr1,3*^f/f^* and S1Pr1,3*^ECKO^* mEC after 4-OHT (1μM, 72h) and siS1PR3 (40nM, 72h) treatments (n≥3/group). LC-MS/MS quantification of total specific **(F-G)** ceramides and **(H-L)** glucosylceramides in Spns2*^f/f^* and Spns2*^ECKO^* mEC after 4-OHT (1μM, 72h) treatment (n=5/group). Data are expressed as mean±SEM. \**P*≤0.05; \*\**P*≤0.01; \*\*\**P*≤0.001. Statistical significance was determined by unpaired t-test.

